# A Powerful Approach for Identification of Differentially Transcribed mRNA Isoforms

**DOI:** 10.1101/002097

**Authors:** Yuan-De Tan, Joel. R. Neilson

## Abstract

Next generation sequencing is being increasingly used for transcriptome-wide analysis of differential gene expression. The primary goal in profiling expression is to identify genes or RNA isoforms differentially expressed between specific conditions. Yet, the next generation sequence-based count data are essentially different from the microarray data that are continuous type, therefore, the statistical methods developed well over the last decades cannot be applicable. For this reason, a variety of new statistical methods based on count data of transcript reads has been correspondingly developed. But currently the transcriptomic count data coming only from a few replicate libraries have high technical noise and small sample size bias, performances of these new methods are not desirable. We here developed a new statistical method specifically applicable to small sample count data called mBeta t-test for identifying differentially expressed gene or isoforms on the basis of the Beta t-test. The results obtained from simulated and real data showed that the mBeta t-test method significantly outperformed the existing statistical methods in all given scenarios. Findings of our method were validated by qRT-PCR experiments. The mBeta t-test method significantly reduced true false discoveries in differentially expressed genes or isoforms so that it had high work efficiencies in all given scenarios. In addition, the mBeta t-test method showed high stability in performance of statistical analysis and in estimation of FDR. These strongly suggests that our mBeta t-test method would offer us a creditable and reliable result of statistical analysis in practice.

---

Development of high-throughput sequencing technologies in recent years (Cloonan et al. 2008a; Cloonan et al. 2008b; Mortazavi et al. 2008) has massively been increasing genomic data and led sequencing cost to rapidly go down so that the sequencing technologies as platforms for studying gene expression or sub-gene expression have become more and more attractive (McCarthy et al. 2012). Current next generation sequencing (NGS) technologies such as RNA-seq (Cloonan et al. 2008a; Cloonan et al. 2008b; Mortazavi et al. 2008), Tag-seq (Morrissy et al. 2009), deepSAGE (t Hoen et al. 2008), SAGE-seq (Wu et al. 2010), and PAS-seq (Shepard et al. 2011) can generate short reads of sequence tags, that is, sequences of 35-300 bp that correspond to fragments of the original RNA. In particular, 3P- seq or PAS-seq (Shepard et al. 2011), a deep sequencing-based method for quantitative and global analysis of RNA polyadenylation has been used to study expression behavior of RNA isoforms in a variety of human and mouse cells.

To evaluate differential expression between conditions or cases, sequences need to be mapped to genome and annotated. After doing so, the sequence data can be transformed to count data at genomic level of interest. Although RNA-Seq can be used to study differential transcription of novel exons, splice-variants and isoforms-specific (Denoeud et al. 2008; Li et al. 2010; Pan et al. 2008; Wang et al. 2009) and allele-specific expression (Degner et al. 2009; Montgomery et al. 2010), our focus here is on differential expression of genes or isoforms due to alterative polyadenylation signals and cleavage sites in 3’ untranslated regions (3’UTR).

A RNA sample may be thought of as a RNA population and each RNA sequence as one individual. Sequencing a RNA sequence is a random process of sampling from a RNA population. If each individual RNA has equal chance to be selected for sequencing, then probability of sequencing a RNA sequence is proportional to the length of waiting time (Anders et al. 2010). Thus number of RNA read counts for a given genomic feature should follow Poisson distribution. The Poisson distribution implicates that only one parameter determines count variation of reads. However, since the Poisson model is just with respect to noise but does not deal with biological source of variation, that is, difference in transcription between samples, some counts are over-dispersed between samples under the Poisson Model. Accordingly, variation of read counts consists of two parts: noise (Poisson) and biological variability, that is, 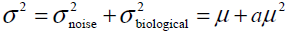 where 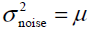 is variance of Poisson distribution and 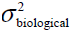 is biological variance, which is determined by biological effect *a*. This is just characteristic of negative binomial distribution (Anders et al. 2010; Robinson et al. 2008).

To identify differential transcription of RNA tags, many statistical methods have been proposed so far. At early stage, most of methods are based on Poisson distribution (Madden et al. 1997) or normal approximations (Kal et al. 1999; Man et al. 2000; Michiels et al. 1999), permutation (Zhang et al. 1997), beta distribution (Baggerly et al. 2003) or over-dispersed logistic linear distribution (Baggerly et al. 2004). As RNA count data have become more and more prevalent, newer statistical methods such as Exact test (Robinson et al. 2008), empirical Bayesian method (Hardcastle et al. 2010), DESeq (Anders et al. 2010), generalized linear modeling (McCarthy et al. 2012), and likelihood ratio tests (Wang et al. 2010) have recently been developed.

Despite the development of technologies decreasing costs of sequencing, RNA-Seq experiments still remain expensive for many researchers so that RNA-Seq studies have to be limited to only a very small number of replicate libraries for each condition or case. The basic scientific need to assess differential transcription due to biological variation remains undiminished but the problem becomes complicated by the fact that different genes or transcripts may have different degrees of biological variation. There is therefore a need to estimate biological variation as reliably as possible from a few replicate libraries (McCarthy et al. 2012). The classical statistical methods are not applicable for such data. To address this problem, many methods such as empirical Bayesian method (baySeq) (Hardcastle et al. 2010), DESeq (Anders et al. 2010) and Exact test (Robinson et al. 2010b; Robinson et al. 2007; Robinson et al. 2008) adopt variation information across transcriptome and the GLM (McCarthy et al. 2012)uses similarity information between genes. However, none of these methods considers fudge effect of small sample size. So-called fudge effect is such an effect on which differences between two means are small but sample variances are also much smaller such that statistics are inflated (Tusher et al. 2001, Tan et al. 2007). This is because sample size is so small that there is a big chance to give rise to small difference among replicates in a large-scale data (see Discussion Section for more detail). The fudge phenomenon broadly exists in high-throughput data, especially, in transcriptomic data because there are a lot of very small counts. Therefore, to suppress such an effect can greatly improve performance of the statistical methods in identification of differentially expressed genes or isoforms. For doing so, we are required to develop novel methods or to modify the existing statistical methods.

Our development work is based on Beta t-test of Baggerly et al (2003) (Baggerly et al. 2003) because this method is not sensitive to data distributions (see Discussion Section). On the other hand, the Beta t-test approach optimizes weights for replicate libraries. The weighting and optimal strategy may be useful for excluding artificial or technical noise in count data and hence the genes or isoforms with better consistent counts in replicates libraries but having differential transcriptions between conditions would be identified with higher probabilities. The third, a very important point, is that it is t-test, a classical distance-variance test approach, that is clear and simple to understand gene differential expression but its fudge effect is also significant. For this reason, we are highly motivated to develop a novel beta t-statistic by which gene mRNAs to be tested can be separated into two different groups with least type I and type II errors.

## Results

### Statistical Methods

Here we follow the notations of Baggerly et al (Baggerly et al. 2003) but for the convenience of description, we use isoforms as features of study. However, our method is available to all types of count data. Let *X*_*i*_ be count for an isoform of interest in library *i*. Let *p*_*i*_ be true proportion of an isoform and *N*_*i*_ be total count in library *i*, that is, size of library *i*. We suppose that the proportion *p*_*i*_ of a count in library *i* follows a beta distribution,

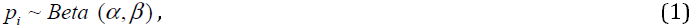

but as mentioned above, the count for an isoform has binomial or negative binomial distribution. In our current study, we consider the binomial distribution instead of negative binomial distribution (see Discussion Section). Since *p̂*_*i*_ = *X*_*i*_/*N*_*i*_ is an estimate of *p*_*i*_, the mean and variance of the estimated proportion for this isoform in library *i* are given by *α*, *β* and *N*_*i*_ (see Supplemental Appendix A).

Considering the case of small sample size, we use weight to correct biases of expectation and variance of estimated proportion p. Supposing that we have *m* replicate libraries in a condition, the mean and variance of proportions in *m* replicate libraries can be linearly combined by weights (Baggerly et al. 2003):

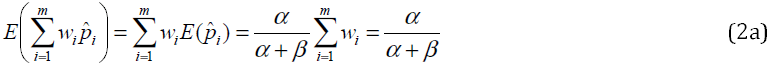

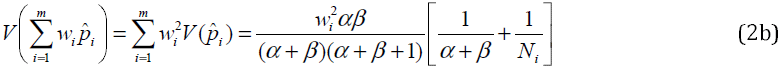

where the sum of weights over *m* replicates is constrained to be 1. Equation (2a) indicates that this combination has the correct mean. Using a partial derivative of variance of weighted proportions with respect to weights, solution for weight vectors can be analytically given by

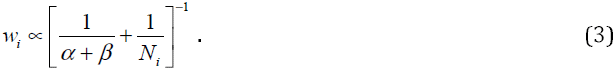

Equation (3) indicates that the weights are determined by the means and sizes of libraries. Here two extreme cases may occur: If *α* + *β* → ∞, then weight *w*_*i*_ is proportional to size of library *i*, *N*_*i*_, meaning that distribution of *p*_*i*_ is degenerate so that there is no change in the true proportion going from sample to sample. If, on the other hand, *α* + *β* is very small, then the weights would be roughly the same for all libraries. The true optimum lies somewhat in between. With the weights, the proportion for an isoform count in a condition is now estimated by

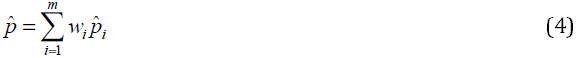

and its variance is also estimated, in an unbiased fashion, by

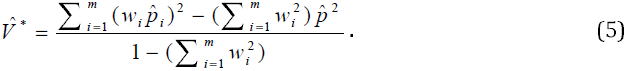

Since we have weights for all parameters (*α*, *β*, *p*, and *V*) in a condition, then an iterative search algorithm for optimal estimation of these parameters can be driven by weights (see Supplemental Appendix B).

Despite the estimation of variances of proportions in a condition is unbiased and optimized, those isoforms of extremely small counts would have very small and similar proportions in a few replicate libraries, which leads variances to be much smaller than differences between means so that the t-values are inflated (see Discussion Section). To avoid occurrence of this phenomenon, a modified estimator of variance is found to be

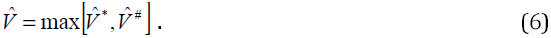

In Baggerly et al (Baggerly et al. 2003), *V̂*^#^ is given by

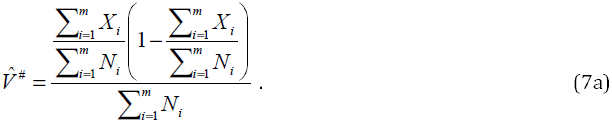

From Equation (7a), we can find that when 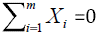, then *V̂*^#^ = 0. In addition, 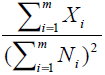 allows *V̂*^#^ to be extremely small. In order to avoid the extreme small variance, we modify *V̂*^#^ as

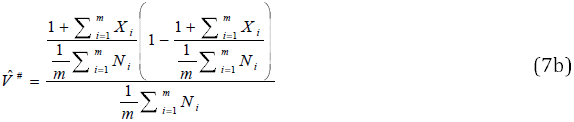

*V̂*^#^ in Equation (7b) is larger than that in Equation (7a). Equation (7b) shows that (1) lower bound of *V̂*^#^ is 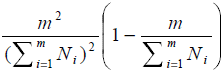, not zero and (2) *V̂*^#^ > *V̂*^*^ is when 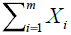 is extremely small. Thus *V̂* = *V̂*^#^ in case of extremely small *V̂*^*^.

By the above optimal estimation, we obtain *p̂*_*A*_ and *p̂*_*B*_, *V̂*_*A*_ and *V̂*_*B*_ in conditions A and B, respectively. Using these estimates, the t-statistic similar to Z statistic suggested by Kar et al (Kal et al. 1999) is found to be

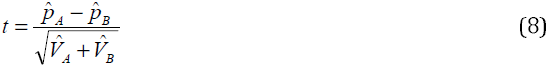

with degrees of freedom

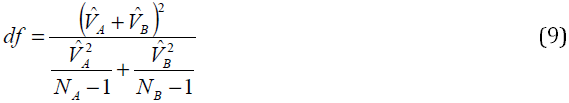

(Baggerly et al. 2003) where 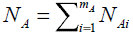 and 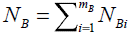 With *df*, we can obtain p-value for each t-statistic from the t-distribution. For count-based transcriptional data, however, since replicate numbers are very small (for example, 3 replicate libraries), a potential for “fudge” effect exists in a population. Although Equation (6) inserts another estimator of variance as a lower bound so as to avoid occurrence of zero or extremely small variance, the fudge effect still exists in Equation (8) due to small sample size. To remove this effect, the t-statistic is modified as

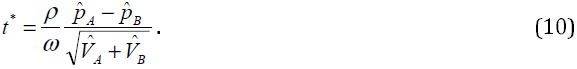

Here *ρ* is defined as 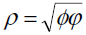 where *ψ* is referred to as “polar ratio” (see Appendix C1).

Equation (C1) indicates that if two data sets *X*_*A*_ = {*X*_*A*1_,…, *X*_*Am*_*A*__)and *X*_*B*_ = {*X*_*B*1_,…,*X*_*Bm_B_*_) for an isoform do not overlap, then *ψ* > 1, otherwise, *ψ* ≤ 1. In statistical theory, two count sets that are definitely separated have a higher probability of showing that they come from two different populations than those that are overlap, *ζ* is referred to as log odds ratio (see Supplemental Appendix C2). If two data sets *X*_*A*_ = {*X*_*A*1_,⋯,*X*_*Am_A_*_) and *X*_*B*_ = {*X*_*B*1_,⋯,*X*_*Bm_B_*_) do not overlap and have a big gap, then *X̄*_*A*_ < *X̄* < *X̄*_*B*_ or *X̄*_*A*_ > *X̄* > *X̄*_*B*_ but 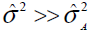 and 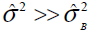, leading to 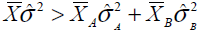 and *ζ* > 1, otherwise, *ζ* ≤ 1 where *X̄* and σ̂^2^ are respectively overall estimated mean and variance of counts for a given isoform; *X̄*_*i*_ and 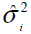 are estimated mean and variance of counts for this isoform in condition *i*, *i* = 1 for A and 2 for B. For *ψ* >1 and *ζ* > 1, we have *ρ* > 1. The t^*^-statistic is potentially preferable to the t-statistic in two aspects: (1) isoforms with small counts are not easily found to have differential expression and (2) t-values with *ρ* > *ω* are inflated but those with *ρ* < *ω* are shrunken. As a result, the fudge effect is suppressed and truly differentially expressed isoforms would be found with a high probability (very low p-value). *ω* is a threshold whose value is inversely proportional to sample size and determined by simulated null data based on the real data. Here we simulate null data, perform our modified Beta t-test(mBeta t-test) with setting *ρ* = 1 and *ω* = 1 for all isoforms on the null simulated data, find false DE isoforms, calculate their *ρ* values, then order them from the smallest to the largest, *ρ*_1_ < *ρ*_2_ <⋯ *ρ*_*j*_ <⋯ < *ρ*_*k*_, and calculate quantiles. We set *q*_1_ = 1/*k*,*q*_2_ = 2/*k*,⋯,*q*_*j*_ = *j*/*k*,⋯*q*_*k*_ = 1 where k is number of false discoveries in a null simulated data. Setting *q*_*j*_ ≥ 0.85, then we choose *ω* = *ρ*_*j*_ value. This means that 85% false discoveries would have *ρ* ≤ *ω* and be excluded. If, however, k ≤5, then *q*_*j*_ ≥ 0.85 is not informative. In this case, we take *ω* = *ρ*_1_. This process is done on all given simulated null datasets. We choose 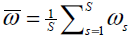 over S simulated null datasets. The *p*-value for each t^*^-value can be obtained from t-distribution using degrees of freedom given or by performing the bootstrap (Storey et al. 2005) (see Supplemental Appendix D).

### Simulation comparisons

We used the 12 scenario stimulation datasets (see Simulation in Materials and Methods) to compare 6 statistical methods for identifying isoforms differentially expressed between two given conditions. The 6 statistical methods chosen here are Beta t-test (Baggerly et al. 2003), empirical Bayesian method (Hardcastle et al. 2010), Exact test (Robinson et al. 2010b; Robinson et al. 2007; Robinson et al. 2008), GLM (McCarthy et al. 2012), DESeq (Anders et al. 2010) and our new Beta t-test (mBeta t-test). The empirical Bayesian (eBayesian) method was implemented on R package baySeq and the Exact test and GLM methods on R package edgeR (Robinson et al. 2010a). DESeq was implemented by R package DESeq (Anders et al. 2010). The Beta and mBeta t-test methods were performed in Matlab. We simulated null data to determine *ω* value for mBeta t-test. We set FDR cutoff = 0.05 and chose estimated FDRs smaller than but closest to this cutoff as acceptable levels for differential expressions of isoforms because the FDR cutoff of 0.05 is widely accepted in multiple tests, especially, in genome-wide studies. We counted isoforms identified to be differentially expressed by these methods, false discoveries and calculated means and standard deviations (stdev) of the numbers of findings and true FDRs for each method chosen and then summarized them in Tables 1–3.

**Table 1.**
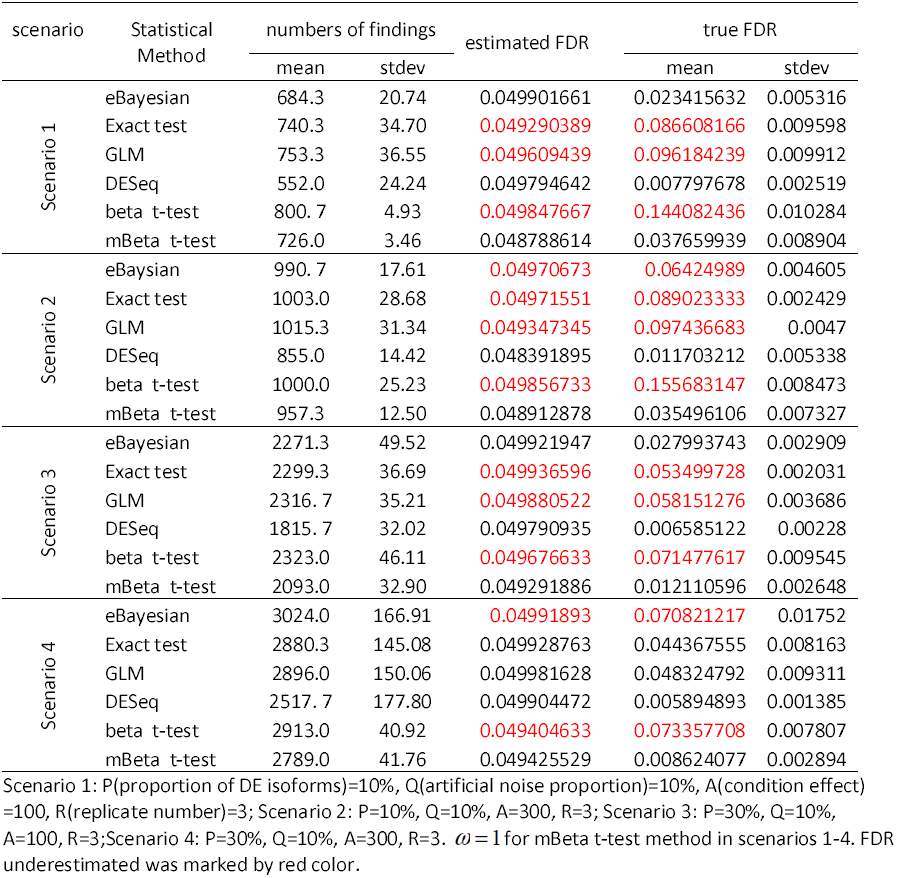
Results of performing statistical methods on simulated data of 18162 isoforms and two conditions in simulation scenarios 1-4 where simulations were based on real transcriptomic data

**Table 2.**
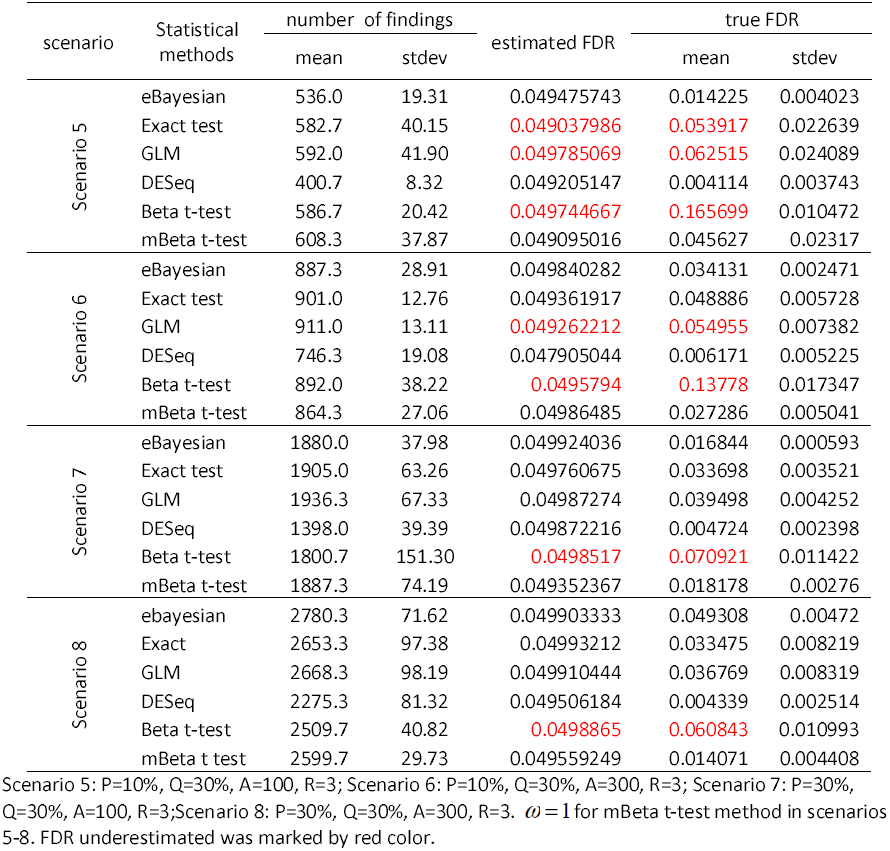
Results of performing statistical methods on simulated data of 18162 isoforms and two conditions in scenarios 5-8 where simulations were based on real transcriptomic data

| scenario | Statistical methods | number of findings |  | estimated FDR | true FDR |  |
| --- | --- | --- | --- | --- | --- | --- |
|  |  | mean | stdev |  | mean | stdev |
| Scenario 5 | eBayesian | 536.0 | 19.31 | 0.049475743 | 0.014225 | 0.004023 |
|  | Exact test | 582.7 | 40.15 | 0.049037986 | 0.053917 | 0.022639 |
|  | GLM | 592.0 | 41.90 | 0.049785069 | 0.062515 | 0.024089 |
|  | DESeq | 400.7 | 8.32 | 0.049205147 | 0.004114 | 0.003743 |
|  | Beta t-test | 586.7 | 20.42 | 0.049744667 | 0.165699 | 0.010472 |
|  | mBeta t-test | 608.3 | 37.87 | 0.049095016 | 0.045627 | 0.02317 |
| Scenario 6 | eBayesian | 887.3 | 28.91 | 0.049840282 | 0.034131 | 0.002471 |
|  | Exact test | 901.0 | 12.76 | 0.049361917 | 0.048886 | 0.005728 |
|  | GLM | 911.0 | 13.11 | 0.049262212 | 0.054955 | 0.007382 |
|  | DESeq | 746.3 | 19.08 | 0.047905044 | 0.006171 | 0.005225 |
|  | Beta t-test | 892.0 | 38.22 | 0.0495794 | 0.13778 | 0.017347 |
|  | mBeta t-test | 864.3 | 27.06 | 0.04986485 | 0.027286 | 0.005041 |
| Scenario 7 | eBayesian | 1880.0 | 37.98 | 0.049924036 | 0.016844 | 0.000593 |
|  | Exact test | 1905.0 | 63.26 | 0.049760675 | 0.033698 | 0.003521 |
|  | GLM | 1936.3 | 67.33 | 0.04987274 | 0.039498 | 0.004252 |
|  | DESeq | 1398.0 | 39.39 | 0.049872216 | 0.004724 | 0.002398 |
|  | Beta t-test | 1800.7 | 151.30 | 0.0498517 | 0.070921 | 0.011422 |
|  | mBeta t-test | 1887.3 | 74.19 | 0.049352367 | 0.018178 | 0.00276 |
| Scenario 8 | ebayesian | 2780.3 | 71.62 | 0.049903333 | 0.049308 | 0.00472 |
|  | Exact | 2653.3 | 97.38 | 0.04993212 | 0.033475 | 0.008219 |
|  | GLM | 2668.3 | 98.19 | 0.049910444 | 0.036769 | 0.008319 |
|  | DESeq | 2275.3 | 81.32 | 0.049506184 | 0.004339 | 0.002514 |
|  | Beta t-test | 2509.7 | 40.82 | 0.0498865 | 0.060843 | 0.010993 |
|  | mBeta t test | 2599.7 | 29.73 | 0.049559249 | 0.014071 | 0.004408 |

Scenario 5: P=10%, Q=30%, A=100, R=3; Scenario 6: P=10%, Q=30%, A=300, R=3; Scenario 7: P=30%, Q=30%, A=100, R=3; Scenario 8: P=30%, Q=30%, A=300, R=3. $\omega = 1$ for mBeta t-test method in scenarios 5-8. FDR underestimated was marked by red color.

**Table 3.** Results of performing statistical methods on simulated data of 18162 isoforms and two conditions in scenarios 9-12 where simulations were based on real isoform transcriptomic data

| scenario | Statistical method | number of findings |  | estimated FDR | true FDR |  |
| --- | --- | --- | --- | --- | --- | --- |
|  |  | mean | stdev |  | mean | stdev |
| Scenario 9 | eBayesian | 1466.7 | 136.99 | 0.049816998 | 0.065909 | 0.017935 |
|  | Exact test | 1455.7 | 135.50 | 0.04942623 | 0.075021 | 0.023157 |
|  | GLM | 1492.0 | 157.43 | 0.049120039 | 0.092843 | 0.032046 |
|  | DESeq | 1270.7 | 116.81 | 0.04968407 | 0.016749 | 0.004976 |
|  | Beta t-test | 1447.7 | 103.58 | 0.049672 | 0.116815 | 0.028366 |
|  | mBeta t-test | 1457.3 | 114.02 | 0.048886388 | 0.03596 | 0.009967 |
| Scenario 10 | eBayesian | 1822.3 | 95.61 | 0.049864672 | 0.104564 | 0.013078 |
|  | Exact test | 1768.3 | 117.50 | 0.049594284 | 0.08398 | 0.028075 |
|  | GLM | 1794.7 | 131.76 | 0.049606781 | 0.094059 | 0.033016 |
|  | DESeq | 1593.3 | 125.52 | 0.049260009 | 0.021667 | 0.012233 |
|  | Beta t-test | 1772.0 | 67.55 | 0.049848 | 0.132159 | 0.025635 |
|  | mBeta t-test | 1697.667 | 67.26 | 0.049301841 | 0.032401 | 0.007489 |
| Scenario 11 | eBayesian | 4894.7 | 289.89 | 0.049930905 | 0.081428 | 0.011789 |
|  | Exact test | 4633.3 | 303.32 | 0.049761768 | 0.051046 | 0.013995 |
|  | GLM | 4681.3 | 332.24 | 0.049960589 | 0.057404 | 0.018121 |
|  | DESeq | 4206.3 | 317.16 | 0.049814792 | 0.013886 | 0.005984 |
|  | Beta t-test | 4700.7 | 362.29 | 0.0493765 | 0.070815 | 0.014574 |
|  | mBeta t-test | 4438.3 | 149.79 | 0.04952921 | 0.0124 | 0.000406 |
| Scenario 12 | ebayesian | 5701.0 | 206.72 | 0.049939398 | 0.099753 | 0.01504 |
|  | Exact test | 5453.7 | 100.48 | 0.049737062 | 0.048492 | 0.006854 |
|  | GLM | 5454.7 | 145.40 | 0.049840291 | 0.053679 | 0.008784 |
|  | DESeq | 4915.3 | 228.00 | 0.049430541 | 0.012464 | 0.005037 |
|  | Beta t-test | 5091.7 | 128.48 | 0.049957167 | 0.066915 | 0.013236 |
|  | mBeta t-test | 5072.3 | 135.57 | 0.049633984 | 0.01087 | 0.002098 |

Scenario 9: P=10%, Q=10%, A=100, R=5; Scenario 10: P=10%, Q=10%, A=300, R=5; Scenario 11: P=30%, Q=10%, A=100, R=5; Scenario 12: P=30%, Q=10%, A=300, R=5. $\omega = 0.45$ for the mBeta t-test method in scenarios 9-12. FDR underestimated was marked by red color.

For small condition effect (A=100) or low artificial noise proportion (Q=10%), or low proportion (P=10%) of DE isoforms, the Beta t-test method had higher power. Since mBeta t-test is a modified beta t-test, in the case of low P or small A, mBeta t-test, eBaysian, Exact test and GLM had similar powers, while in higher P and larger A scenarios, Beta and mBeta t-test had lower powers than eBayesian, Exact test and GLM. In all 12 scenarios (Tables 1-3), DESeq always had very low powers. This is because DESeq always had extremely low true FDRs, indicating that DESeq is a very conservative method that would miss many true differentially expressed isoforms in practice. Beta t-test had much higher true FDRs than its estimated FDRs in all these scenarios, meaning that in the Beta t-test method’s findings, there would be much more false discoveries than estimated, so this is not a conservative method. eBayesian showed higher powers in low artificial noise proportion (Tables 1 and 3) but it also had higher true FDRs than estimated FDRs in most cases. In high artificial noise proportion (Q=30%) scenario, eBayesian performed well (Table 2). GLM showed high powers in all 12 given scenarios but its true FDRs were much larger than estimated in 9 scenarios (Table 1–3), suggesting that this method is also not conservative. In low DE isoform proportion (P=10), low artificial noise proportion (Q=10%) or small condition effect (A=100) scenario, Exact test performed poorly because its true FDRs were much larger than its estimated values in most cases, however, in high P(30%), high Q(30%), and large A(300) scenarios, Exact test had a good performance. Similarly to DESeq, mBeta t-test also had lower true FDRs than its estimated values in all 12 given scenarios but its powers were much higher than DESeq (Tables 1-3), showing that the mBeta t-test method is conservative and powerful.

Stability is an important property of a statistical method. To rate stabilities of these statistical methods in performance, we here used standard deviations (stdev) of finding numbers and true FDRs listed in Tables 1-3 as criterion. Small stdev means that this method has a small fluctuation and hence a high stability in identification of DE isoforms, while larger stdev indicates that it has a bigger fluctuation and hence lower stability. Thus, for each scenario, we ordered these methods by using stdev from the smallest to the largest, assigned order scores (from 1 to 6) to them and averaged their order scores over 12 scenarios. Thus, the order score can be used to measure relative stability of a method: the smaller order score, the higher stability. Table 4 summarizes the results of stability analysis. For findings, mBeta t-test had the highest stability, while GLM had the lowest. Beta t-test, eBayesian, DESeq and Exact test got similar order scores and so they had proximate stabilities. For true FDR, as expected, DESeq showed the highest stability. mBeta t-test was in the second highest rank. GLM and Beta t-test were lowest. Exact test and eBayesian showed similar stabilities.

**Table 4.** Stability analysis of methods

| Standard deviation of findings |  | Standard deviation of true FDR |  |
| --- | --- | --- | --- |
| method | Averaged order score | method | Averaged order score |
| mBeta t-test | 2.333333 | DESeq | 1.833333 |
| Exact test | 3.166667 | mBeta t-test | 2.5 |
| Beta t-test | 3.25 | eBayesian | 3 |
| eBayesian | 3.416667 | Exact test | 3.5 |
| DESeq | 3.833333 | Beta t-test | 5.083333 |
| GLM | 5 | GLM | 5.083333 |

To comprehensively rate these statistical methods, we follow Tan (Tan 2011) and define work efficiency of a statistical method as

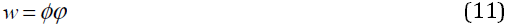

where 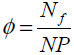 and *φ* = 1 if true FDR<=0.05 and *φ* = 0, otherwise. Here *N* is number of isoforms in simulated data, P, proportion of DE ioforms given in simulation, *N*_*P*_ = *NP*, and *N*_*f*_, number of isoforms found to be differentially expressed by a statistical method. *ϕ* is used to measure power (ability or probability) of a statistical method for identifying a differentially expressed isoform and *φ* to measure conservativeness of this method. The performance of a method must be evaluated by its power and conservativeness. If a method has a high power to find DE isoforms with no conservativeness, its findings would then be unreliable and incredible; if a method has a low power with high conservativeness, then the method would loss many findings. So such two types of statistical methods would have low work efficiencies in identification of DE isoforms.

Table 5 lists work efficiencies of the 6 large-scale statistical methods in 4 pairs of scenarios. From Table 5, one can find that eBayesian and GLM had higher work efficiencies in 3 replicate libraries than in 5 replicate libraries. This is because in 5 replicate libraries, the two methods underestimated FDR at cutoff α=0.05 (Tables 1–3) so that they lost conservativeness. Beta t-test had work efficiency of zero in all scenarios. Exact test had lower efficiencies in low P, low Q, small A and 3 replicates than in high P, high Q, large A and 5 replicates. For the DESeq and mBeta t-test approaches, the work efficiency was greatly raised with increment of sample size. This is due to the fact that the two methods promoted its power with the same conservativeness. Similar results also can be seen in condition effects A=100 versus A=300. These two methods did not significantly respond to change in proportion of DE isoforms and in artificial noise proportion. However, in all simulated scenarios, the mBeta t-test method showed the highest work efficiencies.

**Table 5.** Averaged work efficiencies of statistical methods

| Scenario factors | eBayesian |  | Exact test |  | GLM |  | DESeq |  | Beta t-test |  | mBeta t-test |  |
| --- | --- | --- | --- | --- | --- | --- | --- | --- | --- | --- | --- | --- |
|  | mean | stdev | mean | stdev | mean | stdev | mean | stdev | mean | stdev | mean | stdev |
| 3 replicate libraries | 0.48697 | 0.321 | 0.3726 | 0.40689 | 0.27557 | 0.3884 | 0.57566 | 0.15142 | 0 | 0 | 0.69207 | 0.12025 |
| 5 replicate libraries | 0 | 0 | 0.46259 | 0.53764 | 0 | 0 | 0.81272 | 0.15142 | 0 | 0 | 0.87067 | 0.07198 |
| proportion of DE genes/isoforms=10% | 0.30974 | 0.35327 | 0.13241 | 0.32434 | 0 | 0 | 0.63812 | 0.09414 | 0 | 0 | 0.75333 | 0.14388 |
| proportion of DE genes/isoforms=30% | 0.3281 | 0.36791 | 0.49808 | 0.42951 | 0.16917 | 0.31848 | 0.66774 | 0.18908 | 0 | 0 | 0.7625 | 0.14959 |
| artificial effect proportion =10% | 0.31775 | 0.36784 | 0.21165 | 0.42329 | 0.2128 | 0.42559 | 0.62857 | 0.19842 | 0 | 0 | 0.7298 | 0.1188 |
| artificial effect proportion =30% | 0.3281 | 0.36791 | 0.49808 | 0.42951 | 0.16917 | 0.31848 | 0.66774 | 0.13814 | 0 | 0 | 0.7625 | 0.14959 |
| condition effect =100 | 0.3827 | 0.30323 | 0.23505 | 0.37554 | 0.09485 | 0.23234 | 0.5427 | 0.19842 | 0 | 0 | 0.66057 | 0.12078 |
| condition effect =300 | 0.2666 | 0.41316 | 0.57015 | 0.44847 | 0.27258 | 0.4228 | 0.76666 | 0.1635 | 0 | 0 | 0.84263 | 0.07676 |

In order to display FDR profiles along FDR cutoff, we here plotted averaged true FDRs against averaged estimated FDRs by these compared methods from cutoff = ~ 0 to ~ 0.21 in scenario 1. To evaluate the FDR profiles of these statistical methods, we also plotted a theoretical FDR profile (a diagonal line for true FDR against true FDR) for each method. Figure 1 shows that the estimated FDR curves of eBayesian, GLM and Exact test and Beta t-test are much below their theoretical lines, indicating that these methods, especially, Beta t-test, indeed heavily underestimated their FDRs, while DESeq too much overestimated its FDRs along the cutoff. Therefore, DESeq indeed is a too stringent and too conservative method. The mBeta t-test method overestimated FDR and hence is conservative.

**Figure 1.**
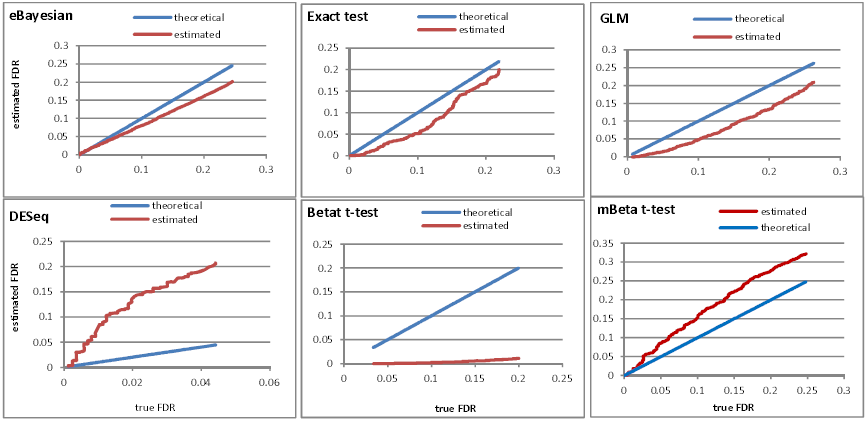
Profiles of true versus estimated FDR. Estimated curve was made by plotting estimated FDR against true FDR along cutoff of ~0 to ~ 0.21 and theoretical line is a diagonal line made by plotting true FDR against true FDR along the same cutoff. The true FDR was calculated by counting false positives in findings of a statistical method at an FDR cutoff point in a simulated dataset and the estimated FDR was predicted by a statistical method. The true and estimated FDRs given figure 1 were averaged over three simulated datasets in scenario 1.

### Real experimental data

The simulated data are generally made in a known distribution and all factors impacting on differential expressions are well controlled, hence evaluation of these statistical methods on the simulated data is conducted in ideal and known states. Obviously, such an evaluation has a limited significance for their application. Real data are therefore required in comparison of statistical methods. However, since everything, in particular, noise distribution in real data is unknown, a direct evaluation of statistical methods is impossibly realized by comparison between true and estimated FDRs. For this reason, we here employed an indirect way for doing so. First, we compared the performances of these statistical methods on our two real datasets: Jurkat T-cell isoform transcriptomic data and Jurkat T-cell gene transcriptomic data. The Jurkat T-cell isoform dataset contains 64428 isoforms which attribute to 14603 genes. These 64428 isoforms were annotated according to alternative poly(A) and cleavage sites within genes and studied on differential transcriptions between resting and stimulating immune states each with 3 replicate libraries by using PAS-seq (Shepard et al. 2011). We used edgeR (Robinson et al. 2010a) to normalize the Jurkat T-cell isoform and gene data. After filtering, 13409 isoforms in the isoform data and 9572 genes in the gene data were available for differential expression analyses. eBayesian had no results because either it was running too long (over at least 2 days in isoform data, infinite loop might occur) or showed NA result (in gene data). GLM obtained 4376 genes and 5039 isoforms of being differentially expressed from our gene and isoform data, respectively, at FDR cutoff of 0.05. Since ratios of findings are so high (45% in the gene data and 37% in the isoform data), we did not believe that this method could work on these real data. DESeq found 261 DE genes and 287 DE isoforms at the same FDR cutoff. The ratios of findings are so low (3% in gene data and 2% in isoform data), basically, DESeq also did not work on the two transcriptomic data. The Beta t-test, mBeta t-test and Exact test methods worked and the results obtained at FDR cutoff of 0.05 are listed in Table 6.

**Table 6.** Results of performances of 3 statistical methods on two real gene transcriptomic data

| data type |  | Exact test | mBeta t-test<sup>a</sup> | Beta t-test |
| --- | --- | --- | --- | --- |
| Gene count data | # of DE genes found | 799 | 1774 | 2515 |
|  | estimated FDR | 0.0499 | 0.0499 | 0.0489 |
|  | least true FDR | 0.2753 | 0.0056 | 0.3062 |
| Isoform count data | # of DE isoforms found | 1029 | 1981 | 3025 |
|  | estimated FDR | 0.0499 | 0.0498 | 0.0499 |
|  | least true FDR | 0.2799 | 0.0101 | 0.3530 |

a: $\omega=1$

In the next step, we compared their findings using Venn Diagram Generator (http://simbioinf.com/mcbc/applications/genevenn/). Figure 2A shows that except that the three methods shared 554 DE genes, Exact test and mBeta t-test shared 22 DE genes in their 2019 findings while the former and Beta t-test shared merely 3 DE genes in their 2760 findings. Similar result was also found in the isoform data (Fig. 2B). In addition, if a gene was found to be differentially expressed by a method only, then it is highly possible that this DE gene would be falsely discovered. Figure 3 visualizes heat maps of 10 genes identified to be differentially expressed by mBeta t-test method (Fig. 3A), 770 by Beta t-test only (Fig. 3B), and 220 by Exact test only (Fig. 3c). Indeed, the method-specific genes do not display obvious expression difference between no stimulation (NS) and 48h poststimulation. Thus, we defined no share ratio of findings (*m*_*i*_/*M*_*i*_ where *m*_*i*_ is number of method *i*-specific findings, and *M*_*i*_ is numbers of findings identified by method *i*) as least true false discovery rate (least true FDR is corresponding to q-value defined by Storey et (Storey et al. 2003)). Using this indirect method, we obtained the least true FDRs for the findings of the Exact test, mBeta t-test and Beta t-test methods, respectively, in the two real transcriptomic datasets (Table 6). From Table 6, one can see that Exact test and Beta t-test extremely underestimated their FDRs while mBeta t-test still overestimated FDR in its findings. These are well consistent with those found in the simulated data.

**Figure 2.**
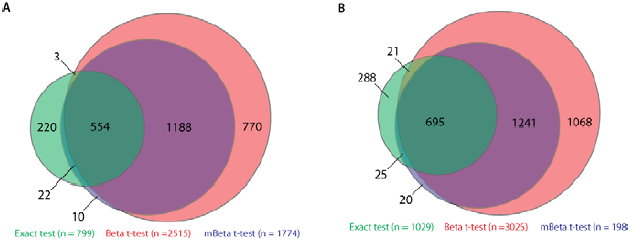
Venn diagram analyses ofthe Exact test, Beta t-test and mBeta t-test approaches. A: DE genes found in Jurkat T-cell gene transcriptomic data. B: DE isoforms found in Jurkat T-cell isoform transcriptomic data.

**Figure 3.**
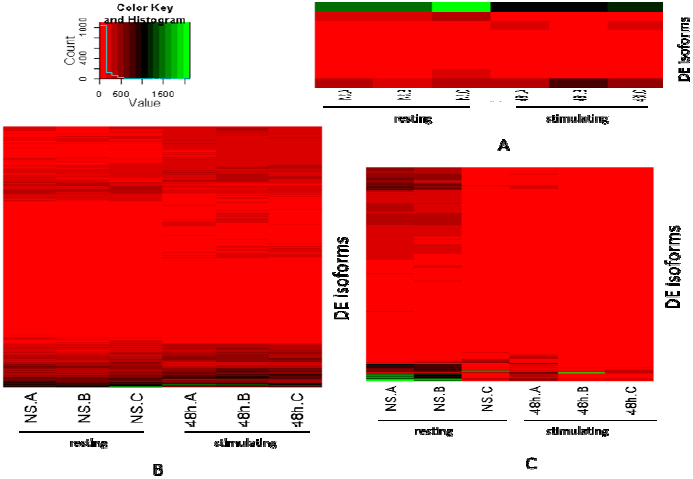
Heatmaps of method-specific differentially expressed genes. NS.A, NS.B, NS.C are replicate libraries A, B and c in resting state (no stimulation). 48h.A, 48h.B and 48h.c are replicate libraries A, B and c in stimulating state (48h poststimulation). Method-specific DE genes were treated as false DE genes. Heatmaps show that these genes really do not have obvious expression difference between resting and stimulating states. A: 10 DE genes found by mBeta t-test only. B: 770 DE genes found by Beta test only. C: 220 DE genes found by Exact test only.

### Experimental Validation

qRT-PCR experiments were carried out to valid our method’s findings. Our qRT-PCR experiments were done in Jurkat T-cells at rest and 48 hours after stimulation. To make our qRT-PCR results be more representative, we randomly chose the genes that were found to be up-regulated or down-regulated at 48 hours after stimulation and not to be differentially expressed by mBeta t-test. We used relative differences between stimulation and rest and relative variation coefficient (VC) (see Materials and Methods Section) to do comparison between the RNA-seq and qRT-PCR data. The results show that genes UBL3, MST123 and KIAA0465 that were found to be up-regulated to respond to stimulation (blue columns in Fig. 4A) in RNA-seq data also displayed positive response to stimulation in qRT-PCR data (red columns in Fig. 4A). Gene CD47 negatively responded to stimulation in both datasets while gene TESK2 was not detected to have significantly difference between stimulation and rest in these two datasets. In expression direction and relative expression amount, these two datasets show cc=0.9 (Pearson correlation coefficient) (Fig. 4A), suggesting that our transcriptomic data were well consistent with qRT-PCR data. Using relative VC, we found that gene UBL3 had small expression noise at rest and stimulation in these two datasets, while genes TESK2 had bigger expression noises in the transcriptomic data (Fig. 4B). This is why gene UBL3 was detected to be differentially expressed but TESK2 was not though both UBL3 and TESK2 had small counts of mRNA reads in transcriptomic data.

**Figure 4.**
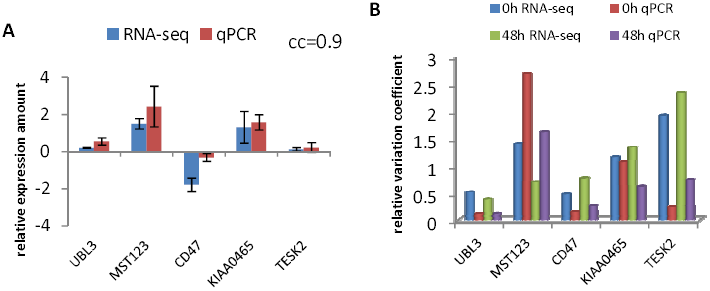
Comparison between RNA-seq data and qRT-PCR data on 5 genes chosen. In RNA-seq data genes UBL3, MST123, CD47 and KIAA0465 were found to be differentially expressed between rest and stimulation. TESK2 had no differential expression. Genes UBL3, MST123, KIAA0465 were significantly higher at stimulation than at rest, while genes CD47 and BCL11B were significantly lower at stimulation than at rest. A: Relative expression comparisons of 5 genes between RNA-seq and qRT-PCR. In RNA-Seq data, relative expression of a gene is defined as *d*_*g*_ / *d̄* where *d*_*g*_ = *n̄*_*gt*_ – *n̄*_*g*0_, *d̄* is averaged value, g= UBL3, MST123, CD47, KIAA0465 and TESK2, *n̄* is averaged count of reads over three replicates and t = 48 hour. In qRT-PCR, the relative expression of a gene is defined as Δ_*g*_ – Δ _*b*_ where 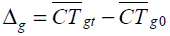, *b* = background gene for control, and 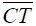 is averaged CT value over three replicates and CT is log2 transformed threshold value of amplification in qRT-PCR. In our experiment, we used TBP (TATA binding protein) as background expression because it has no change in expression with time. B: Relative expression variation coefficient comparison of 5 genes between RNA-seq and qPCR data. Relative expression variation coefficient is defined as 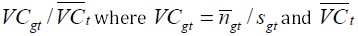 is averaged variation coefficient over all selected genes at time t of stimulation.

## Discussion

Although our simulated data were made in the NB distributions, Beta and mBeta t-test worked well if estimation of FDR was not considered. This is because not only the NB counts can be well approximated by binomial distributions but also the frequencies (p) of counts of mRNA reads in libraries can be approximated by beta distributions. While the eBayesian, GLM, Exact test, and DESeq approaches are merely based on the NB distribution. Therefore, for real datasets whose distributions are often unknown, mBeta t-test will perform well. For example, as seen in Result Section, eBayesian and GLM, DESeq could not work on our Jurkat T-cell isoform and gene data, while Exact test, Beta t-test and mBeta t-test worked even though their results have big differences. We also applied these methods to our leukemia transcriptomic data containing 10299 genes (data not yet published), the results show that all methods chosen can work but eBayesian has very low power (it just found 165 DE genes), while GLM, Exact test and mBeta t-test identified, respectively, 780, 733 and 711 DE genes and hence performed very similarly. These strongly suggest that eBayesian and GLM are specific to the NB distribution.

In addition, in genome-wide data, especially, in transcriptomic data, sample sizes usually are very small, for example, 3 or 2 replicate libraries in each condition due to high cost and biological resource limitation. Small samples would lead to a fudge effect (Tan 2011; Tan et al. 2011; Tan et al. 2006). For example, in count data containing more than ten thousands of isoforms, two 3-replicate small-count sets would have a larger probability of showing that they would be sampled from the same distribution not only than two 5-replicate small-count sets but also than two 3-replicate big-count sets. On the other hand, in transcriptome-wide data, small-count data have more chance to be weakly fluctuated by noise and to form extremely small within-variances than big-count data, giving rise to inflating t-statistics. For general statistical methods, the genes or isoforms with small count data in small samples would easily be found to be differentially expressed between conditions due to inflation of statistic. To address this problem, many methods developed for identifying differentially expressed genes in microarray data introduce a constant to shrink statistics. For example, In SAM (Tusher et al. 2001), two-sample t-test is modified as S-test by adding a minimized coefficient of variation S_0_ of differences between two conditions to denominator. In the regularized *t* test (Baldi et al. 2001), two-sample t-test is modified by combining gene-specific variance with global average variance. The two methods shrink all t-statistics across the whole genes detected in microarrays. So they have low powers (Tan 2011). Tan et al (Tan et al. 2006) used a conditional shrinking method to address the problem of inflating t-test. But this conditional shrinking method cannot be introduced to Beta t-test because the Beta t-test is based on differences in frequencies (proportions) of tags between conditions (Baggerly et al. 2003).

Baggerly et al (Baggerly et al. 2003) employed a weighting and iteration strategy to look for an optimal estimation of parameters beta and alpha of frequency that is assumed to follow beta distribution for a tag in a condition and furthermore developed a new t-test, we called Beta t-test. Weight and optimization is a strategy for excluding artificial or technical noise in count data. Although Baggerly et al (Baggerly et al. 2003) recognized small-count effect on t-tests and tried to avoid the problem of t-value inflation using alternative variance given in Equation (18a), our practice demonstrated that Equation (18a) does not substantially reduce the fudge effect. For this reason, we modify the alternative variances by utilizing means of total counts over all libraries in a condition for those isoforms with very small counts. Analytically, it can be seen that the alternative variance defined in Equation (18b) is larger than that in Equation (18a). Our simulation really showed that the above small-count effect on testing for differential expressions of isoforms was mostly reduced by our modified alternative variances.

In order to eliminate small sample effect, we introduced a gene-or isoform-specific variable *ρ* into the Baggerly et al.s’ (Baggerly et al. 2003)Beta t-test. *ρ* is used to measure overlap between two count sets. If two count sets more overlap, then *ρ* becomes smaller; if two count sets separate, then *ρ* > 1. The larger gap between two count sets, the larger *ρ*. In theory, two count sets that are separated have a higher probability of showing that they came from two different populations than those that overlap. Besides, if noise within count sets is large, then *ζ* is small, which makes *ρ* become small. Thus, *ρ* shrinks t-values of overlapped count sets and inflates t-values of separated count sets with small noise. As seen in Tables 1-3, compared to the Beta t-test method, our mBeta t-test approach did not obviously decrease its power but significantly reduce false discovery rate so that it has higher work efficiencies. Considering sample size effect, we set a threshold *ω* for *ρ*. That is, t-values are inflated with *ρ* > *ω*, or shrunken with *ρ* < *ω* As a result, almost all of small t-values are compressed into a short interval close to zero but the t-values with *ρ* > *ω* are further enlarged and a region in which truly differential expressions of isoforms and expression noises are mixedly distributed becomes very narrow (Fig. 5). Since p-value only depends on t-value given degree of freedom, p-values with inflating t-values become smaller while those with shrinking t-values become larger. Thus a lot of false discoveries are also compressed into the zero neighbors so that very few false positives would be found (Fig. 5). Threshold *ω* depends on sample size. The larger sample size, the smaller *ω*. However, when sample size is large, *ω* becomes very small, ability of *ρ* controlling false discoveries becomes very weak because the gaps between two datasets are vanished and there is not fudge effect in such data.

**Figure 5.**
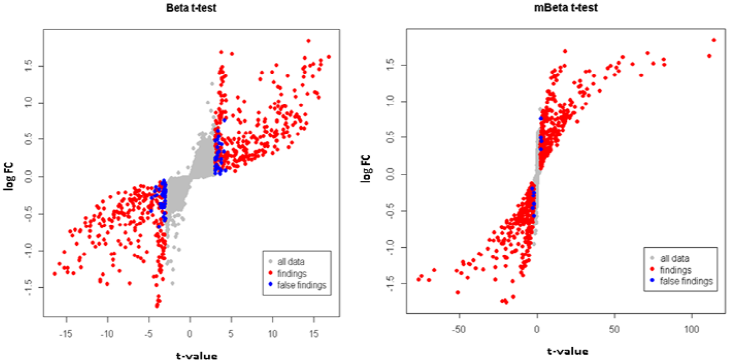
Plots of t-statistics versus log FC. logFC=log (mean in 48h poststimulation/mean in NS) and t-values were given by two beta t-test methods from the simulated data in which 10% of isoforms randomly assigned with condition effect *τ* ≤ 100 were differentially expressed between two conditions each with three replicate libraries. Simulation was conducted by NB pseudorandom generator based on real isoform transcriptomic data. A: The Beta t-statistics are distributed in interval between -16 and 16, while those for no differential expressed genes in Beta t-test are distributed in interval between −4.8 and 4.8. Thus a lot of false discoveries (blue dots) are distributed in the neighboring areas of t-value ≥4.8 or t-value ≤ −4.8. B: Plots of the mBeta t-statistics versus log FC. The t-statistic interval is enlarged from below −50 to over 100, while the t-statistics for no differential expressed genes are strongly compressed into a very narrow area close to zero. Thus a lot of false discoveries of the Beta t-test method are also moved into this area so that very few false positives (blue dots) would be found.

ROC is popularly used to evaluate statistical methods (Hardcastle et al. 2010) (Robinson et al. 2007) but ROC has two fatal drawbacks. First, ROC cannot evaluate the FDR estimation. For multiple tests, since FDRs are unknown, they must be estimated so that one can determine which features have statistical significances. Precise or conservative estimation of FDRs is important for an experimental scientist or statistician to choose a statistical method in practice because if a method significantly underestimates FDRs in findings, then it would provide much more false findings than expected or if it much overestimates FDRs, we would then loss many true findings (Tan 2011). Second, in some cases, the ROC curves of the methods are tightly close to each other or overlap together, meaning that these methods have similar sensitivity against specificity but it does not suggest that they have the same or approximate performances because they may have different estimations of FDRs. For example, we employed simulated data of scenarios 1 and 4 to makeup eBayesian, Exact test, DESeq and mBeta t-test ROC curves. The results show that mBeta t-test performed best at specificity <0.3 in scenario 1 (Fig. 6A) or <0.5 in scenario 4 (Fig. 6B), DESeq had the slight higher sensitivity than mBeta t-test when specificity > 0.3 in scenario 1 (Fig. 6A) or >0.5 in scenario 4 (Fig. 6B), eBayesian had the lowest curve, Exact test and GLM had almost the same curve, performed better than eBayesian when specificity >0.4 (Fig. 6). However, these ROC curves did not show significant difference, in particularly, in scenario 4. For this reason, our evaluation of these methods chosen was based on comparison between true and estimated FDRs. The simulated data showed that the eBayesian and GLM methods worked well in the ideal NB distributions, lower proportions of DE isoforms, smaller condition effect and smaller number of replicate libraries but they performed poorly in the case in which sample sizes were larger and proportion of DE isoform was higher or condition effect was bigger. This is because in such a scenario they generally had a high power to find DE isoforms but, on the other hand, their FDRs are often notably underestimated. Therefore, we evaluated performance of a statistical method in power (ability or probability to find DE genes or isoforms) and conservativeness of FDR estimation (reliability of findings). A statistical method with high power but no conservativeness of FDR estimation would offer us a lot of unreliable findings or a statistical method with low power but conservativeness would miss many true DE genes or isoforms. That is to say, these two types of methods would have low work efficiency to perform their statistical analysis of real data.

**Figure 6.**
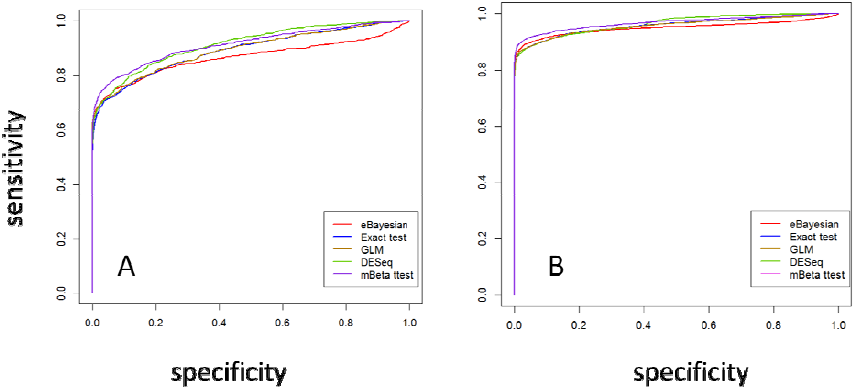
ROC comparison of statistical methods. ROC curves of the eBayesian, Exact test, GLM, DESeq and mBeta t-test methods were made from simulated datasets. Sensitivity = true positive fraction (TPF) and specificity = false positive fraction (FPF). A: simulated data came from scenario 1 (proportion of differentially expressed isoforms = 10%, technical noise proportion = 10% and treatment effect A = 100, sample size = 3) and B: from scenario 4 (proportion of differentially expressed isoforms = 30%, technical noise proportion = 10% and treatment effect A = 300, sample size = 3).

It is required to indicate that the Exact test method performs very fast, in contrast, the eBayesian method (baySeq) is very intensive in computation and took a long time (maybe infinite loop occurred in calculation of posterior probabilities when performed on our real data). Nevertheless, the current version of baySeq is not available for our real data. The mBeta t-test method is also computationally intensive because it runs literately to look for an optimal estimation of weight and beta and alpha parameters for each isoform. However, it finishes its work in 15 minutes for more 10 thousands of isoforms if we don’t use bootstrap to calculate p-values.

## Materials and Methods

### Cell Lines and Stimulation

Jurkat T-cell lines were obtained from the ATCC and maintained in RPMI (ATCC) with 10% fetal bovine serum supplemented with penicillin and streptomycin (Gibco). Jurkat T-cells were stimulated with plate-bound antibodies coated with a solution of 1 ⟆g anti-CD3 (OKT3 - eBiosciences) and 5 ⟆g anti-CD28 (CD28.2 - BD Pharmingen). Activation of T-cell was monitored via flow cytometry detection of CD69 expression (FN-50) 48 hours after stimulation (Simms et al. 1996).

### High-throughput Sequencing Library Generation

Total RNA was harvested from resting and stimulated cells with Trizol reagent and processed as per manufacturer instructions. Polyadenylated RNA was isolated with the Poly(A)-Purist MAG kit as per manufacturer instructions. Libraries for high-throughput sequencing were established as described (Shepard et al. 2011) with minor modifications. Libraries were sequenced via 50 bp paired-end sequencing on an Illumina GAIIx in genome sequencing center in Baylor College of Medicine. The listed reagents were from Life Technologies.

### Annotation and pipeline analyses

Paired-end reads were first subjected to a profiler removing A and T homopolymer runs within the forward and reverse reads, respectively. Pairs in which the length of both reads was greater than 25 bp were mapped to the human genome reference (UCSC hg19) with Bowtie(Langmead et al. 2009) using default parameters. The reads were simultaneously mapped to the UCSC KnownGene database to identify putative exon-spanning reads or pairs. The union of mate pairs mapping uniquely to the genome and those mapping specifically to the KnownGene database were condensed to isoforms by 3’ alignment identity, filtered for false priming, and assigned gene identity and region (e.g. CDS, 3’ UTR) using UCSC KnownGene annotations. Cleavage and polyadenylation sites were defined as the median genome coordinate of all reads within a 20-base pair sliding window of an adjacent read. Defined sites were carried forward for analysis only if they were present in all libraries representing an individual cellular type or state. Intralibrary isoform representation was normalized to pseudocounts using the negative binomial method(Robinson et al. 2008).

### qRT-PCR

Total RNA was respectively isolated from Jurkat T-cells at rest and 48hours after stimulation using TRIZOL (Invitrogen). RNA was digested with DNase I to remove contaminating genomic DNA. One microgram of total RNA was used to generate cDNA with the ImProm-II™ Reverse Transcription System (Promega) and real-time PCR was performed in triplicates in an Eppendorf Mastercycler. qRT-PCR was performed using a pair of primers (forward and reverse primers) designed for a selected gene. ΔΔC_T_ method was used to calculate relative quantitation of qRT-PCR product on a 7500 Fast Real-Time PCR machine with SDS software (Applied Biosystems).

### Simulations

To evaluate statistical properties of various approaches, we used the NB pseudorandom generator to create RNA isoform count datasets in 12 scenarios, each repeated three times for calculations of means and standard deviations. Our simulations were conducted on our Jurkat T-cell isoform data from which we took 18290 isoforms and 3 replicated libraries in each of two conditions (resting and stimulating states). We set two levels (A=100 and 300) of condition (or treatment) effect on differential transcription of isoforms and linearly assigned the effects *τ* = UA to differentially expressed isoforms where U is uniform variable (*U* ∈ (0,1]), two levels of proportions of differentially expressed isoforms: P =10 and 30%, two levels of artificial noise proportions: Q=10 and 30% and 2 levels of sample sizes: R=3 and 5 replicate libraries. Here artificial noise (also called technique or Poisson noise) indicates that the noise does not comes from experimental system error but come from techniques such as sequencing, mapping, annotation and pipline analysis, etc. In simulated data, isoforms with averaged count <5 were filtered, thus 18162 isoforms were available for analysis.

### Software

A package for implementing mBeta t-test was written in Matlab and a program for generating simulation data was written in R. They are available for request.

## Acknowledgement

We are indebted to Natee Kongchan for library preparation and Alexander Ruch and Jixin Deng for primary processing of our data sets. This work was funded by The Cancer Prevention and Research Institute of Texas (HIRP100475) and the National Cancer Institute (CA126752, CA131474).

